# Improved functions for non-linear sequence comparison using SEEKR

**DOI:** 10.1101/2024.03.10.584286

**Authors:** Shuang Li, Quinn Eberhard, Luke Ni, J. Mauro Calabrese

## Abstract

SEquence Evaluation through *k*-mer Representation (SEEKR) is a method of sequence comparison that utilizes sequence substrings called *k*-mers to quantify non-linear similarity between nucleic acid species. We describe the development of new functions within SEEKR that enable end-users to estimate p-values that ascribe statistical significance to SEEKR-derived similarities as well as visualize different aspects of *k*-mer similarity. We apply the new functions to identify chromatin-enriched long noncoding RNAs (lncRNAs) that harbor *XIST*-like sequence fragments and show that several of these fragments are bound by *XIST*-associated proteins. We also highlight the best practice of using RNA-Seq data to evaluate support for lncRNA annotations prior to their in-depth study in cell types of interest.

## Introduction

Mammalian genomes produce thousands of long noncoding RNAs (Mattick, Amaral et al. 2023). While the majority remain functionally uncharacterized, lncRNAs have been shown to carry out a diversity of molecular functions, which can be specified by their sequence and structural properties, or even merely through the act of their transcription. However, relative to most protein coding RNAs, lncRNAs are poorly conserved, evolve rapidly, and rarely harbor long stretches of linear sequence similarity. As a result, identifying the sequence elements that give rise to molecular functions in lncRNAs remains a challenge (Mattick, Amaral et al. 2023).

In 2018, we reported the development of a simple algorithm to perform non-linear sequence comparison, called SEquence Evaluation through *k*-mer Representation (SEEKR). In SEEKR, sequences of interest are compared by their standardized *k*-mer contents, where a *k*-mer is defined as a short sequence substring of length *k*. Using SEEKR, we showed that *k*-mer content can be used to identify similarities in protein-binding and localization between lncRNAs, and to identify lncRNA sequences that may share molecular functions. The main outputs of SEEKR are lists of standardized *k*-mer contents within the sequences being compared (i.e. their “*k*-mer profiles”), along with matrices of Pearson’s r values that quantify similarity in *k*-mer profiles between sequences. These outputs can help to identify features shared within lncRNAs of interest, even if those features are not detectable by linear alignment (Kirk, Kim et al. 2018, Sprague, Waters et al. 2019, Kirk, Sprague et al. 2021).

However, in its current form, SEEKR lacks a framework to assess the significance of *k*-mer similarity relative to a background expectation. In contrast, the most broadly used linear alignment algorithms present end-users with p-values that describe the significance of similarity between pairs of sequences (e.g. BLAST; (Altschul, Gish et al. 1990)). These p-values enable rapid parsing of results and prioritization of similarities that may be biologically significant. Herein, we describe the implementation of a p-value function along with other updates that improve the interpretability of *k*-mer similarities defined by SEEKR. As a framework to illustrate these updates, we compare the lncRNA *XIST* to the set of human lncRNAs annotated by GENCODE (Frankish, Carbonell-Sala et al. 2023). We identify several chromatin-associated lncRNAs that contain significant similarity to protein-binding modules within *XIST*. These lncRNAs include the known repressive lncRNA *KCNQ1OT1*, the architectural lncRNA *NEAT1*, and other lncRNAs with uncharacterized molecular functions. We also document examples of lncRNAs whose transcript annotations are not strongly supported by short-read RNA-Seq data in specific cell types, highlighting the best-practice of evaluating lncRNA annotations for context-dependent validity before drawing conclusions about the molecular function or biological role of a lncRNA. SEEKR updates are accessible through Github, the Python Package Index, and the Docker Hub.

## Results

### New SEEKR functions including those to estimate significance of *k*-mer similarity

SEEKR estimates the similarity of pairs of lncRNAs by standardizing their length-normalized *k*-mer frequencies relative to those found in a larger, user-specified set of background sequences. In controlled tests, we have found that SEEKR performs best when using a value of *k* where 4^*k* is approximately equal to the average length of the sequences being analyzed including those in the background set, a practice which reduces the number of *k*-mers with count values of zero in any given RNA while maintaining discriminatory power (Kirk, Kim et al. 2018, Sprague, Waters et al. 2019). We have also found it important to select a set of background sequences that is large enough to capture the extent of sequence variation across the transcriptome of the species being analyzed. For example, when comparing the *k*-mer content of the human lncRNA *XIST* to that of another lncRNA, it would be reasonable to use a large collection of spliced human lncRNAs as the set of background sequences, such as the collection annotated by GENCODE (Frankish, Carbonell-Sala et al. 2023), and use *k*-mer values of *k*=4, 5, or 6, because the average length of a lncRNA in the GENCODE collection is on the order of one kilobase (kb).

Additionally, lncRNA collections such as GENCODE often contain partially identical lncRNA sequences that arise from the presence of related transcript isoforms (Frankish, Carbonell-Sala et al. 2023). As a consequence, lncRNAs genes that have large numbers of annotated spliceforms or alternative transcriptional start or end sites may cause certain *k*-mers to be overrepresented in the background distribution as a function of the lncRNA annotation process rather than the biology being studied. Thus, in most cases, a preferred set of background sequences is one that contains a single representative transcript for each lncRNA gene. For example, starting with the GENCODE v43 lncRNA collection, the total number of lncRNA transcripts is reduced from 58,023 to 15,550 by selecting the set of deduplicated “Ensembl_canonical” lncRNAs that are greater than 500nt in length (henceforth referred to as GENCODE canonical lncRNAs).

Once a *k*-mer length and set of background sequences have been selected, lncRNA comparisons can be made. SEEKR reports its similarity scores in the form of a Pearson correlation coefficient, r, which quantifies the level of linear correlation between any two lncRNAs’ standardized *k*-mer contents. Pearson’s r values, along with standardized *k*-mer contents, can be examined between individual pairs of lncRNAs as well as en masse, the latter often in the context of a heatmap or network graph. However, a shortcoming of Pearson’s r is that as a standalone similarity metric, it provides no information about what level of similarity might be expected when comparing two randomly selected sequences. For example, comparing the human lncRNA *XIST* to that of another human lncRNA, *KCNQ1OT1*, using a *k*-mer length of *k*=6 and the GENCODE canonical lncRNAs as a background set, we find that SEEKR returns a Pearson’s r of 0.07. Intuitively, this low Pearson’s r value suggests that relative to the set of canonical lncRNAs, the 6-mer profiles of *XIST* and *KCNQ1OT1* exhibit a weak positive correlation. But it remains unclear what value of r might be expected when comparing any two lncRNAs using this *k*-mer length and this set of background sequences. Quantifying that expectation would enable the parsing of results to consider only those similarities (or differences) that are the most significant.

To meet this need, we developed a series of functions within the SEEKR package that enable end-users to estimate the significance of SEEKR-derived Pearson’s r values. We also created new functions to visualize of different aspects of *k*-mer similarity, and deprecated other functions that were either redundant with new ones or difficult to install using current versions of Python (Figure 1A). Below, we describe the use of these several of these functions in a biological context. Updates to SEEKR are available for download through the Python Package Index, GitHub, and the Docker Hub (https://pypi.org/project/seekr/; https://github.com/CalabreseLab/seekr; https://hub.docker.com/r/calabreselab/seekr). A separate GitHub page contains the code and console commands used to generate the data displayed in the figures below (https://github.com/CalabreseLab/seekr2.0_update_manuscript).

**Figure 1.**
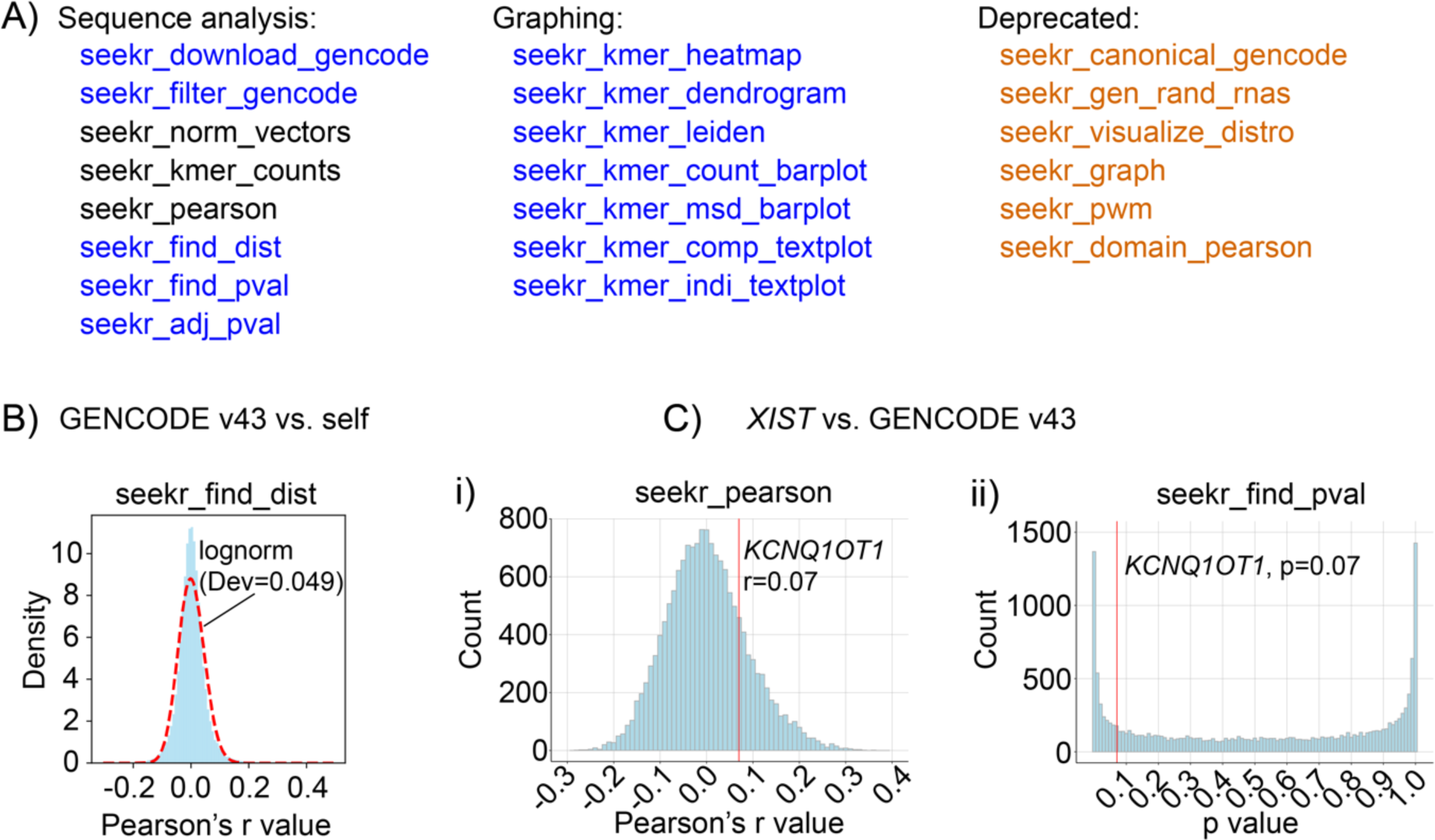
New SEEKR functions including those to estimate significance of *k*-mer similarity. **(A)** Console / command-line functions in SEEKR. Blue text, new or updated functions. Black text, retained functions. Orange text, deprecated functions. “seekr_filter_gencode” replaces “seekr_canonical_gencode”. “seekr_kmer_leiden” replaces “seekr_graph”. The “fasta-shuffle-letters” utility in the MEME Suite (Bailey, Johnson et al. 2015) can be used in place of “seekr_gen_rand_rnas”. **(B)** Density of Pearson’s r values derived from a self vs. self comparison of the non-identical set of GENCODE v43 Ensembl_canonical lncRNAs at *k* = 6, fit to a log-normal distribution. Dev, deviation from a log-normal distribution, calculated as the D statistic of the Kolmogorov-Smirnov test, which quantifies the distance between the empirical distribution function of the sample and the cumulative distribution function of the fitted distribution. **(C)** *XIST* compared to the non-identical set of >500nt long GENCODE v43 Ensembl_canonical lncRNAs at *k* = 6. **(i)** Distribution of Pearson’s r values for all comparisons. **(ii)** Distribution of p values for all comparisons. Red lines mark values for *XIST* vs. *KCNQ1OT1*. The SEEKR console / command line functions that were used to generate data in (B) and (C) are shown above the figure graphs.

To estimate the significance of SEEKR-derived Pearson’s r values, using the console / command line, end-users first employ the “seekr_find_dist” function to evaluate what distribution is the best fit for their set of selected background sequences. In our experience, SEEKR-derived Pearson’s r values for large sets of background sequences are often well-fit by log-normal distributions (Figure 1B). Next, end-users employ their selected distribution in the “seekr_find_pval” function to derive p-values that describe the probability that any pair of lncRNAs are more similar to each other than would be expected from random chance. Finally, if desired, end-users can perform correction for multiple-testing using “seekr_adj_pval”. Continuing from the example above, we find that *XIST* and *KCNQ1OT1* are more similar to each other than ∼93% of sampled pairwise comparisons of GENCODE canonical lncRNAs, despite their low positive Pearson’s r value (p-value of 0.07; Figure 1C). Moreover, with two additional commands, we find 3183 GENCODE canonical lncRNAs whose overall *k*-mer contents are more similar to *XIST* than *KCNQ1OT1* (Table S1). The top ten most *XIST*-similar lncRNAs, along with their Benjamini-Hochberg adjusted p-values are shown in Table 1.

**Table 1.**
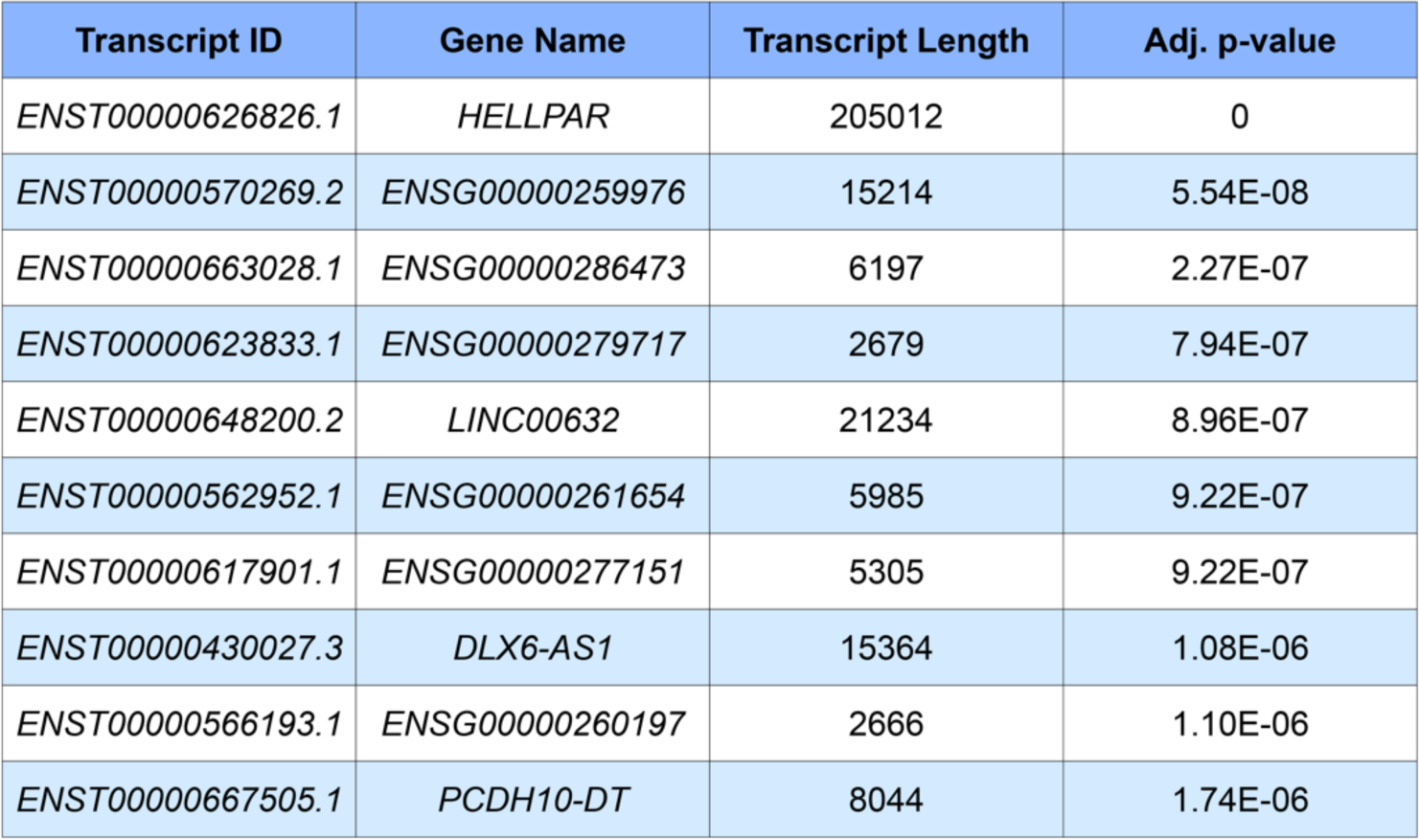
Top ten most *XIST*-similar lncRNAs, as assessed by overall *k*-mer content. We evaluated the set of non-identical “Ensembl_canonical” GENCODE v43 lncRNA transcripts that were ≥500 nt in length. Transcript Length is shown in nucleotides. SEEKR-derived p-values were adjusted using the Benjamini-Hochberg method.

### SEEKR functions to visualize features of *k*-mer similarity

We next applied SEEKR to investigate *LINC00632*, *DLX6-AS1*, and *PCDH10-DT*, which are highly similar to *XIST* (Table 1) and whose Ensembl_canonical transcript isoforms are strongly supported by short-read RNA-Seq data (de Goede, Nachun et al. 2021)*. XIST* is thought to encode its repressive functions through the cumulative action of separate domains that interact with different subsets of proteins (Trotman, Braceros et al. 2021). While full repression by *XIST* requires its entire sequence, some of its regions most critical for repression are comprised of tandemly repeated sequences, referred to as Repeats A, B, D, E, and F (Dixon-McDougall and Brown 2021, Dixon-McDougall and Brown 2022). Therefore, to determine whether our select *XIST*-like lncRNAs harbored any regional similarity with *XIST*, we separated the lncRNAs into ∼500 nucleotide (nt) fragments and compared the fragments in each lncRNA to each fragment within *XIST*. Because data suggest Repeats A, B, D, E, and F function as discrete domains, we evaluated each *XIST* Repeat as a single intact fragment and separated the intervening *XIST* intervals into ∼500 fragments (Files S1 and S2 contain fragments of *XIST* and all other GENCODE canonical lncRNAs used in this study, respectively). From these analyses, we found that *LINC00632*, *DLX6-AS1*, and *PCDH10-DT* each harbor significant similarity to Repeat E but lack similarity to the other Repeats in *XIST* (Figures 2B-D). Moreover, these three lncRNAs also harbor significant similarity to fragments distributed across the final exon of *XIST* (Figures 2B-D). A closer examination revealed that the similarities could be attributed to a uniform enrichment of *k*-mers rich in A and T nucleotides; whereas, *k*-mers rich in G and C nucleotides were among the most variably enriched (Figures 2E-G). We also observed that many of the 500nt fragments within *XIST*’s final exon were significantly more similar to each other than would be expected by chance (Figures 2H, 2I, and S1).

**Figure 2.**
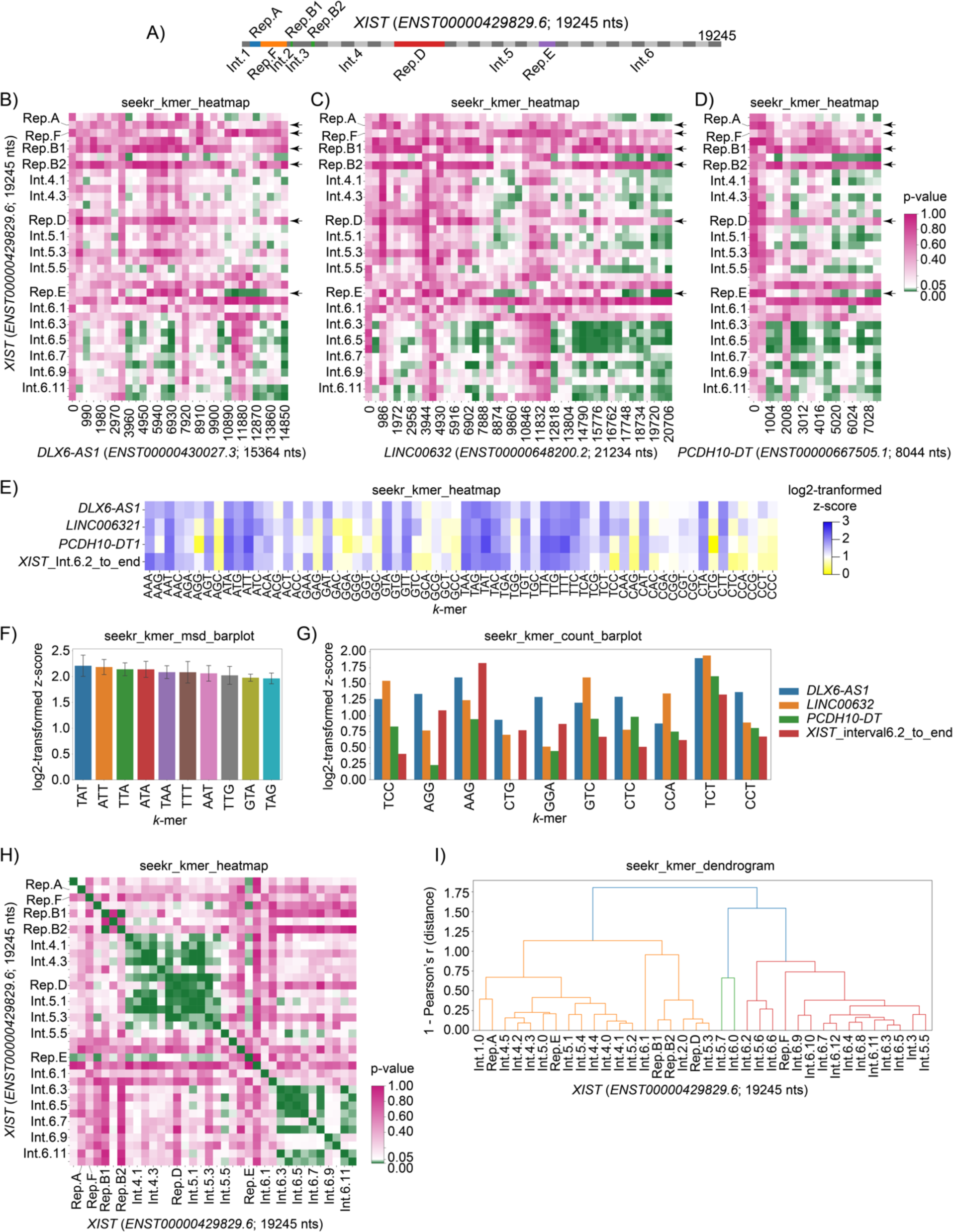
SEEKR functions to visualize features of *k*-mer similarity. **(A)** Intervals / chunks of *XIST* used to identify regional similarities. Repeats A-F in *XIST* are colored in non-gray shades and were each evaluated as single intervals, regardless of length. “Repeat F” is annotated here as a region that spans nts 743 to 1807 in *XIST* (*ENST00000429829.6*). **(B-D)** Regional similarities between *XIST* and *LINC00632*, *DLX6-AS1*, and *PCDH10-DT*. The latter three lncRNAs were separated into ∼500nt consecutive intervals and compared to the *XIST* intervals in (A). Heatmaps show p values for each comparison. Comparisons were made using *k* = 4. **(E)** Heatmap displaying log2-tranformed z-scores at *k* = 3, for the 3′ portion of *XIST* (Interval 6.2 to its transcript end) and *LINC00632*, *DLX6-AS1*, and *PCDH10-DT* relative to the set of non-identical “Ensembl_canonical” GENCODE v43 lncRNA transcripts :≥500 nt in length. Log2-transformed z-scores were derived using seekr_kmer_counts. *k* = 3 was used here (instead of *k* = 4) to increase the legibility of *k*- mer sequences in the figure panel. **(F)** Top ten 3-mers that are the most enriched in lncRNAs from (E) relative to the set of “Ensembl_canonical” also from (E). **(G)** Top ten 3-mers that exhibit the most variable enrichment in lncRNAs from (E) (i.e. highest standard deviation across all four lncRNAs). **(H)** Regional similarities within *XIST*. Heatmap shows p-values for each pairwise comparison of *XIST* intervals shown in (A). Comparisons were made using *k* = 4. **(I)** Dendrogram showing clusters of *XIST* intervals from (A) that harbor related *k*-mer contents. Clusters (different colors) were defined as the nodes whose correlation distances are less than 0.7*[max_distance_in_dendrogram]. Comparisons were made using *k* = 4. The SEEKR console / command line functions that were used to generate graphs are displayed above each figure panel.

### Domain-based search identifies *XIST*-like lncRNAs and highlights variable support among lncRNA annotations

Given that the tandem repeats within *XIST* are some of the regions most essential for its ability to induce and maintain gene silencing, we used SEEKR to perform a parallel search for *XIST*-like lncRNAs among the set of GENCODE canonical lncRNAs. In this search, we separated all 15,550 GENCODE canonical lncRNAs into ∼500nt fragments (including *XIST*, as a positive control), and then used SEEKR to identify lncRNAs that contain fragments with significant *k*-mer similarity to *XIST* Repeats A, B, D, E, and F (unadjusted p-value of <0.05). We then summed the number of *XIST*-similar fragments in each lncRNA and used these sums to rank lncRNAs by their overall *XIST*-likeness. This analysis identified several intriguing lncRNAs (Table 2; Table S2). As might have been expected, *XIST* ranked highly (3^rd^). However, the known repressive lncRNA *KCNQ1OT1* ranked 8^th^ (in a tie with five other transcripts; (Schertzer, Braceros et al. 2019, Quinodoz, Jachowicz et al. 2021)). Moreover, 18 of the top 22 lncRNAs were expressed at detectable levels in K562 cells, and of those 18 lncRNAs, 17 were chromatin-enriched, including the architectural lncRNA *NEAT1* (Table 2; (Dunham, Kundaje et al. 2012, Obuse and Hirose 2023)). Thus, with this simple fragment-based search, we identified several chromatin-enriched lncRNAs that harbor domains that resemble those required for repression by *XIST*, including *KCNQ1OT1* and *NEAT1*. Additional *XIST*-like transcripts of interest include *ENST00000605862.6*, *ENST00000506640.3*, and *ENST00000622550.2*, functionally uncharacterized lncRNAs which are both spliced and expressed in multiple cell types (Dunham, Kundaje et al. 2012, de Goede, Nachun et al. 2021), and *RENO1*, a conserved lncRNA whose depletion in mouse embryonic stem cells causes changes in gene expression, a reduction in chromatin accessibility, and a failure to differentiate properly into neurons (Hezroni, Ben-Tov Perry et al. 2020).

**Table 2.**
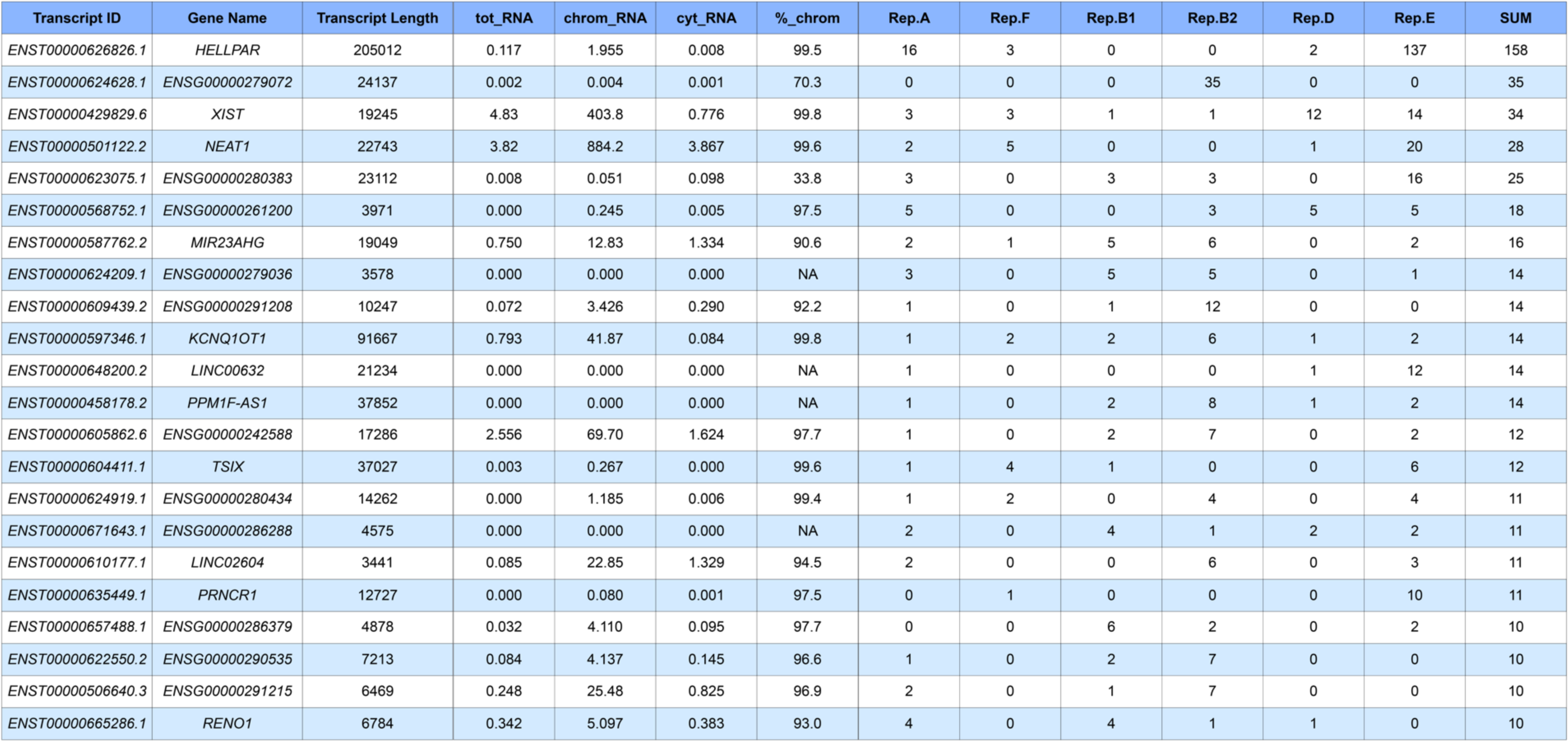
Top 22 *XIST*-like lncRNAs from GENCODE v43. “tot_RNA”, total/ribosome-depletion RNA-Seq data; “chrom_RNA”, chromatin fraction RNA-Seq data; “cyt_RNA”, cytosolic fraction RNA-Seq data; all RNA-Seq data are displayed in TPM units and were collected from K562 cells as part of the ENCODE project (Dunham, Kundaje et al. 2012). “%_chrom”, 100*[chrom_RNA_TPM] / [chrom_RNA_TPM + cyt_RNA_TPM + 0.0001]. “Rep.[A-E]”, count of significant hits to each XIST repeat when analyzing the 500nt chunks of each listed lncRNA. “SUM” sum of counts in “Rep.[A-E]” columns.

The tandem repeats in *XIST* appear in a specific order and are thought to serve as recruitment centers for specific RNA-binding proteins (RBPs (Trotman, Braceros et al. 2021)). With the possible exception of *KCNQ1OT1*, *XIST*-like fragments within the lncRNAs of Table 2 appeared in an order that was different from that found *XIST* (Figure 3). However, we hypothesized that at least some of the fragments might still associate with the same RBPs as their cognate domains within *XIST*. Therefore, in three lncRNAs of interest, we examined the eCLIP read density profiles of six RBPs that have been subject to eCLIP in K562 cells and exhibit characteristic enrichment in the tandem repeats in *XIST* (Van Nostrand, Freese et al. 2020). These RBPs included RBM15 (enriched over Repeat A); HNRNPM (enriched over Repeat F region); HNRNPK (enriched over Repeats B and D); and MATR3, PTBP1, and TIA1 (enriched over Repeat E; Figure 3A). In all three lncRNAs, we observed *XIST*-like fragments that colocalized with eCLIP signal from the expected RBPs. In *KCNQ1OT1*, Repeat A-, M-, and B-like fragments co-localized with RBM15, HNRNPM, and HNRNPK, respectively (Figure 3B). In *NEAT1*, Repeat F-, B/D-, and E-like fragments colocalized with HNRNPM, HNRNPK, and MATR3/PTBP1/TIA1, respectively (Figure 3C). In *ENST00000605862.6*, A- and B-like fragments co-localized with RBM15 and HNRNPK, respectively (Figure 3D). Thus, while the arrangement and number of *XIST*-like fragments in each of those lncRNAs differs from *XIST*, several of their *XIST*-like fragments exhibit enriched association for the expected RBPs. The location of *XIST*-like fragments in all lncRNAs from Table 2 can be visualized relative to their genomic annotations, RNA-Seq, and eCLIP data in the UCSC Genome Browser session associated with this manuscript ((Nassar, Barber et al. 2023); https://genome.ucsc.edu/s/qeberha/seekr_update_3_5_24).

**Figure 3.**
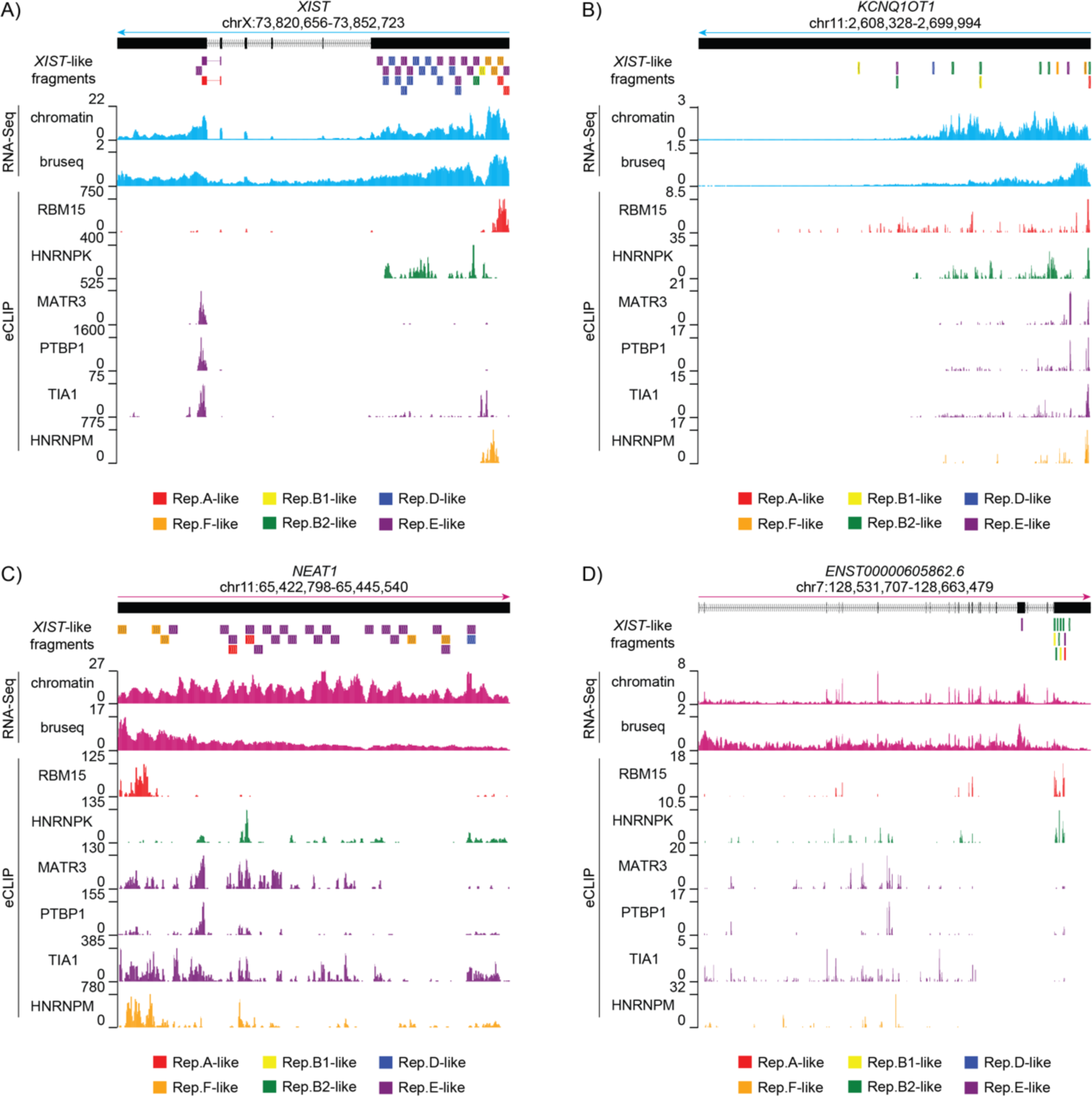
Domain-based search identifies chromatin-associated lncRNAs that harbor *XIST*- like fragments. **(A-D)** Screen images from the UCSC Genome Browser displaying gene annotations, location and identity of *XIST*-like fragments, read density from chromatin-associated RNA-Seq and from Bru-Seq, and background-corrected eCLIP signal for a subset of RNA-binding proteins enriched over the tandem repeats of *XIST*.

During the course of our analyses, we also noted that transcript annotations for several highly ranked lncRNAs in Table 2 were not strongly supported by short-read RNA-Seq data from K562 or HepG2 cells (Dunham, Kundaje et al. 2012). The most highly ranked *XIST*-similar lncRNA, *HELLPAR*, is annotated as an unspliced, 200kb lncRNA that begins at the 3′ end of the protein-coding gene *PARPBP* and terminates near the 5′ end of the protein-coding gene *IGF1* (van Dijk, Thulluru et al. 2012, Frankish, Diekhans et al. 2021). However, RNA-Seq read density suggests that transcription across the *HELLPAR* locus is not due to the expression of a single lncRNA. Rather, it appears to result from a combination of imprecise termination of the upstream *PARPBP* gene, leading to read-through transcription in the 5′ half of the *HELLPAR* locus, and the transcription of a separate lncRNA, *LINC02456*, whose promoter is found in the center of the locus and whose transcribed product runs antisense through the *IGF1* protein-coding gene (Figure 4A; (Dunham, Kundaje et al. 2012, de Goede, Nachun et al. 2021, Zhu, Du et al. 2023)). Likewise, again visualizing short-read RNA-Seq data from (Dunham, Kundaje et al. 2012, de Goede, Nachun et al. 2021), we found examples of other *XIST*-like lncRNAs located in K562 expressed regions but whose transcript annotations were not strongly supported. These included *TSIX*, whose only RNA-Seq read density coincided with the gene structure of *XIST* on the opposite strand (and therefore may have arisen due to imperfect strand-specificity of the dUTP RNA-Seq protocol (Levin, Yassour et al. 2010); Figure 4B); *LINC02604*, which is annotated as a monoexonic lncRNA but appears to be part of a much larger transcribed and spliced region (Figure 4C); and three monoexonic lncRNAs that overlap in the sense direction relative to longer protein-coding or lncRNA genes: *ENST00000623075.1*, *PRNCR1*, and ENST00000624919.1 (Figure 4D-F). While these transcript annotations do harbor *XIST*-like fragments, their annotations are weakly supported by RNA-Seq data, and it could be questioned whether they should be studied as standalone lncRNAs in K562 cells.

**Figure 4.**
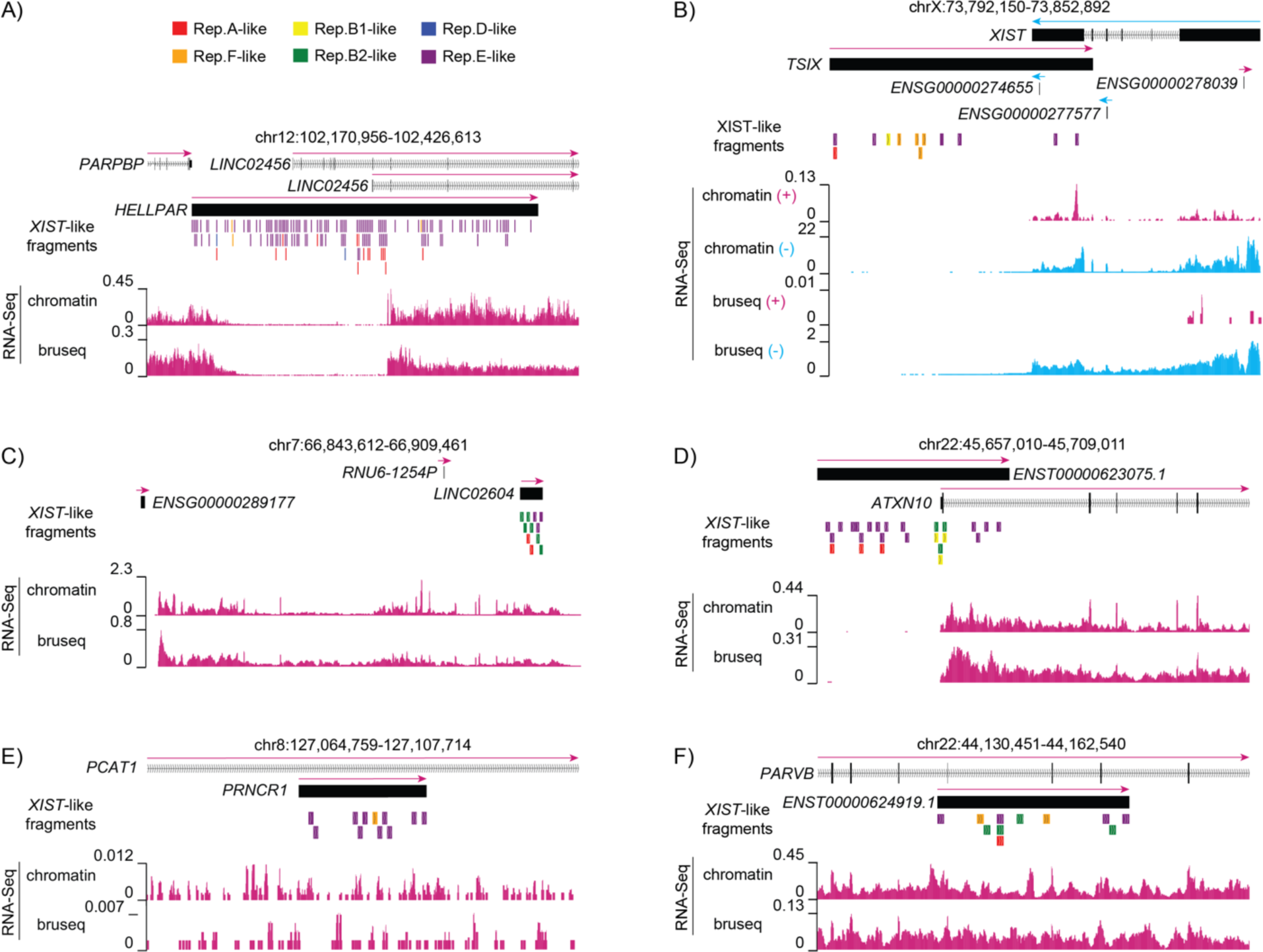
Regions expressed in K562 cells that contain *XIST*-like lncRNA annotations which are not well-supported by RNA-Seq density. **(A-D)** Screen images from the UCSC Genome Browser displaying gene annotations, location and identity of *XIST*-like fragments, and read density from chromatin-associated RNA- and Bru-Seq experiments performed in K562 cells (Dunham, Kundaje et al. 2012, Luo, Hitz et al. 2020).

## Discussion

We describe a series of updates to the SEEKR package that enable enhanced interpretation of SEEKR-derived similarity metrics. The updates were designed for application to lncRNAs but can be applied to the study of any nucleotide sequence. SEEKR is written in python and can be installed via the Python Package Index or the Docker Hub (https://pypi.org/project/seekr/; https://github.com/CalabreseLab/seekr; https://hub.docker.com/r/calabreselab/seekr). The major functions of SEEKR can be implemented via the UNIX console, facilitating their use by biologists with little or no experience with Python. The SEEKR, Python, and UNIX commands used in the analyses above can be found in the GitHub page associated with this manuscript (https://github.com/CalabreseLab/seekr2.0_update_manuscript).

To illustrate the use of the new SEEKR functions, we applied them to the discovery and study of *XIST*-like lncRNAs in the human transcriptome. With a minimal set of commands and python code, we identified several lncRNAs that harbor *XIST*-like sequence features, although none save perhaps *KCNQ1OT1* resembled *XIST* to such an extent that we would predict that the lncRNAs harbored repressive activity. Searches for whole-transcript similarity to *XIST* followed by fragment-based analyses highlighted three lncRNAs – *LINC00632*, *DLX6-AS1*, and *PCDH10-DT* – that contained regions similar to *XIST* Repeat E and its terminal exons, which likely serve architectural roles in *XIST* but may also enable recruitment of certain histone-modifying enzymes (Yamada, Hasegawa et al. 2015, Sunwoo, Colognori et al. 2017, Yue, Ogawa et al. 2017, Pandya-Jones, Markaki et al. 2020, Dixon-McDougall and Brown 2022). We also observed an overall enrichment for A- and T-rich *k*-mers contained within the final long exon of *XIST*.

A fragment-based search for regions similar to the tandem repeats in *XIST* identified a different set of *XIST*-like lncRNAs. Here, the known repressive lncRNA *KCNQ1OT1* ranked 9^th^ among 15,550 lncRNAs queried, and nearly every lncRNA in the top 22 was enriched in the chromatin fraction of K562 cells. Several of the *XIST*-like fragments in these lncRNAs co-localized with expected sets of *XIST*-associated RNA-binding proteins. Our results underscore the utility of fragment-based *k*-mer searches, particularly when the query and target lncRNAs are substantially longer than the presumed functional modules contained within them (Sprague, Waters et al. 2019).

Lastly, our study highlights the best practice of using RNA-Seq data to evaluate lncRNA annotations for support prior to their study in cell types of interest. Visualization of short read RNA-Seq data identified several lncRNA transcripts annotations that are not optimal representations of the lncRNAs produced from those genomic regions in K562 or HepG2 cells. One example was *HELLPAR*, which is annotated as a monoexonic ∼200kb lncRNA. However, RNA-Seq data suggest that the *HELLPAR* locus does not produce a 200kb lncRNA in K562 or HepG2 cells. Instead, the RNA-Seq data are best described by two separate transcripts: one resulting from apparent read-through of an upstream protein-coding gene, and the other resulting from a separately annotated lncRNA whose promoter sits in the center of the *HELLPAR* locus. We found other examples of lncRNAs whose transcript annotations are not strongly supported by RNA-Seq data in K562 or HepG2 cells, nor in many of the tissue types analyzed as part of the GTEx project (Dunham, Kundaje et al. 2012, de Goede, Nachun et al. 2021). It is possible that lncRNAs resembling those annotations are produced in cell types that we did not investigate above. However, the discordance highlights the importance of examining support for transcript annotations before drawing conclusions about a lncRNA in a cell type of interest. Each of the unsupported lncRNAs described in Figure 4 harbored RNA-Seq read density under a portion of their annotations. In these cases, reliance on RNA-Seq count data alone, without visual inspection of the data, would suggest that lncRNA transcripts resembling those annotations were expressed in K562 and HepG2 cells. In the GitHub page associated with this manuscript, we describe how to use custom scripts and standard genomic tools to convert RNA-Seq alignments into wiggle tracks for display in the UCSC Genome Browser (i.e. STAR, Samtools, and BEDtools (Li, Handsaker et al. 2009, Quinlan and Hall 2010, Dobin, Davis et al. 2013, Raney, Dreszer et al. 2014); https://github.com/CalabreseLab/seekr2.0_update_manuscript). We have found that visual inspection of RNA-Seq alignments in this way is a simple yet powerful approach to evaluate the support for lncRNA transcript annotations prior to their experimental characterization.

## Methods

### SEEKR analyses

The Python code and console commands used for *k*-mer analyses in each Figure and Table can be found in https://github.com/CalabreseLab/seekr2.0_update_manuscript. SEEKR updates can be installed via the Python Package Index, via GitHub, or via the Docker Hub (instructions found on https://github.com/CalabreseLab/seekr). Files S1 and S2 contain fragments of *XIST* and all other GENCODE canonical lncRNAs, respectively. The community graph in Figure S1 was made using Gephi (Bastian, Heymann et al. 2009).

### Analysis of ENCODE RNA- and Bru-Seq data

RNA- and Bru-Seq datasets were downloaded from the ENCODE portal (https://www.encodeproject.org/; (Dunham, Kundaje et al. 2012, Luo, Hitz et al. 2020). From K562 cells, the datasets used were: total RNA-seq (ENCSR885DVH), fractionated RNA-Seq (chromatin: ENCSR000CPY; PolyA cytosolic: ENCSR000COK), Bru-Seq (ENCSR729WFH), and Bru-Chase (2hr: ENCSR633UIR; 6hr: ENCSR762OPQ). From HepG2 cells, the datasets used were: total RNA-Seq (ENCSR181ZGR), Bru-Seq (ENCSR974AQD), Bru-Chase (2hr: ENCSR295FEH; 6hr: ENCSR135DZR). Using the STAR aligner, a genomic index was generated using the GRCh38 primary assembly genome FASTA and the GENCODE v43 basic annotation GTF using the ‘--sjdbGTFfile’ flag. Individual replicates from each experiment were then aligned (Dobin, Davis et al. 2013). Alignments were filtered for quality using Samtools view with the ‘-q 30’ flag and the FASTQ sequences for mate pairs R1 and R2 were extracted by ‘samtools fastà, providing the alignments as paired end inputs and the ‘-s’ flag to filter out any singletons (Li, Handsaker et al. 2009). Using kallisto, FASTQ files were aligned to a set of sequences extracted from the GENCODE GRCh38 v43 basic annotation GTF combined with additional transcript sequences that contained all exons and introns from the earliest start to the latest end of each non-monoexonic v43 gene (Bray, Pimentel et al. 2016, Frankish, Carbonell-Sala et al. 2023). Expression values for each GENCODE canonical lncRNA are reported in Table S2.

### Visualization of RNA-Seq data in the UCSC genome browser

Using the quality-filtered STAR alignments from the K562 chromatin fraction RNA-seq (ENCSR000CPY) and K562 Bru-Seq (ENCSR729WFH) datasets, replicates were merged with Samtools using the ‘samtools merge’ command (Quinlan and Hall 2010). To extract negative stranded data from the merged replicate files (both of which were reverse-stranded, paired-ended RNA-Seq experiments), the ‘samtools view -h -f 99’ and ‘samtools view -h -f 147’ commands were used followed by ‘samtools merge’. Conversely, to extract positive stranded data from the merged replicate files, the ‘samtools view -h -f 83’ and ‘samtools view -h -f 163’ were used followed by ‘samtools merge’. These filtered and merged BAM files were converted to BED12 files using BEDtools (Quinlan and Hall 2010). The number of aligned reads were counted from the original filtered and merged alignments using Samtools (Li, Handsaker et al. 2009). A python script (make_wiggle_tracks_1_11_24.py) was then used to generate wiggles from BED12 files. Wiggles were converted into bigWigs using the ucsctools/320 ‘wigToBigWig’ command (Kent, Zweig et al. 2010). See the GitHub page for line-by-line code (https://github.com/CalabreseLab/seekr2.0_update_manuscript).

### Visualization of (s)eCLIP data in the UCSC genome browser

BAM and BED files from (s)eCLIP data experiments (E) and their matched mock input control experiments (C) were downloaded from the ENCODE portal (https://www.encodeproject.org/) for six RNA-binding proteins: RBM15 (E: ENCSR196INN, C: ENCSR454EER); MATR3 (E: ENCSR440SUX, C: ENCSR183FVK); PTBP1 (E: ENCSR981WKN, C: ENCSR445FZX); TIA1 (E: ENCSR057DWB, C: ENCSR356GCJ); HNRNPK (E: ENCSR953ZOA, C: ENCSR143CTS); and HNRNPM (E: ENCSR412NOW, C: ENCSR212ILN) (Dunham, Kundaje et al. 2012, Van Nostrand, Pratt et al. 2016, Luo, Hitz et al. 2020).

Replicates from CLIP experiments were downsampled prior to merging for visualization. The number of aligned reads from each replicate (GRCh38-aligned BAM files) was determined using ‘samtools view -c’ (Li, Handsaker et al. 2009). Replicates were downsampled first by determining the minimum read count between replicates, and then the downsampling_fraction was calculated as follows: [1 – (replicate 1 read counts – replicate 2 read counts) / (minimum read count between the replicates)]. The downsampling_fraction was used in ‘samtools view -b -s <downsampling_fraction>’ to reduce the size of the larger dataset. Downsampled replicates were merged, filtered for MAPQ>30, and split by strand using Samtools (Li, Handsaker et al. 2009). For eCLIP experiments, ‘samtools view -b -q 30 -f 144’ was used to retrieve the second mate of the negative stranded data while ‘samtools view -b -q 30 -f 160’ was used to retrieve the second mate of the positive stranded data. For the seCLIP experiments, ‘samtools view -b -q 30 -f 16’ was used to retrieve the negative stranded data while ‘ samtools view -b -q 30 -F 16’ was used to retrieve the positive stranded data. The ENCODE peak BED files for replicates from each experiment were downloaded and sorted with BEDtools (Quinlan and Hall 2010). For each downsampled and merged eCLIP BAM file, the reads under the peaks from each replicate were extracted using ‘bedtools intersect -split -ubam’ and the resultant BAM files were merged with Samtools (Quinlan and Hall 2010). The reads under the same set of peaks were likewise extracted and merged from the corresponding mock input controls.

Wiggle tracks were then created as described in the RNA-Seq section above. Specifically, peak-extracted BAM files were converted to BED12 files using BEDtools (Quinlan and Hall 2010). The number of aligned reads from the original filtered, downsampled, and merged alignments (pre-peak filtering) were counted with Samtools (Li, Handsaker et al. 2009). A custom python script (make_wiggle_tracks_1_11_24.py) was then used to generate wiggles from BED12 files. Signal for each eCLIP wiggle file was then normalized relative to its corresponding control by subtracting the signal in each bin of the control wiggle from the signal in the same bin of the experiment wiggle. Negative values were excluded. This process was completed with the ‘control_normalize_wiggles_2_20_24.py’ script. Wiggles were then converted into bigWigs using ucsctools/320 ‘wigToBigWig’ command (Kent, Zweig et al. 2010). See the GitHub page for line-by-line code (https://github.com/CalabreseLab/seekr2.0_update_manuscript).

---

**Figure S1.**
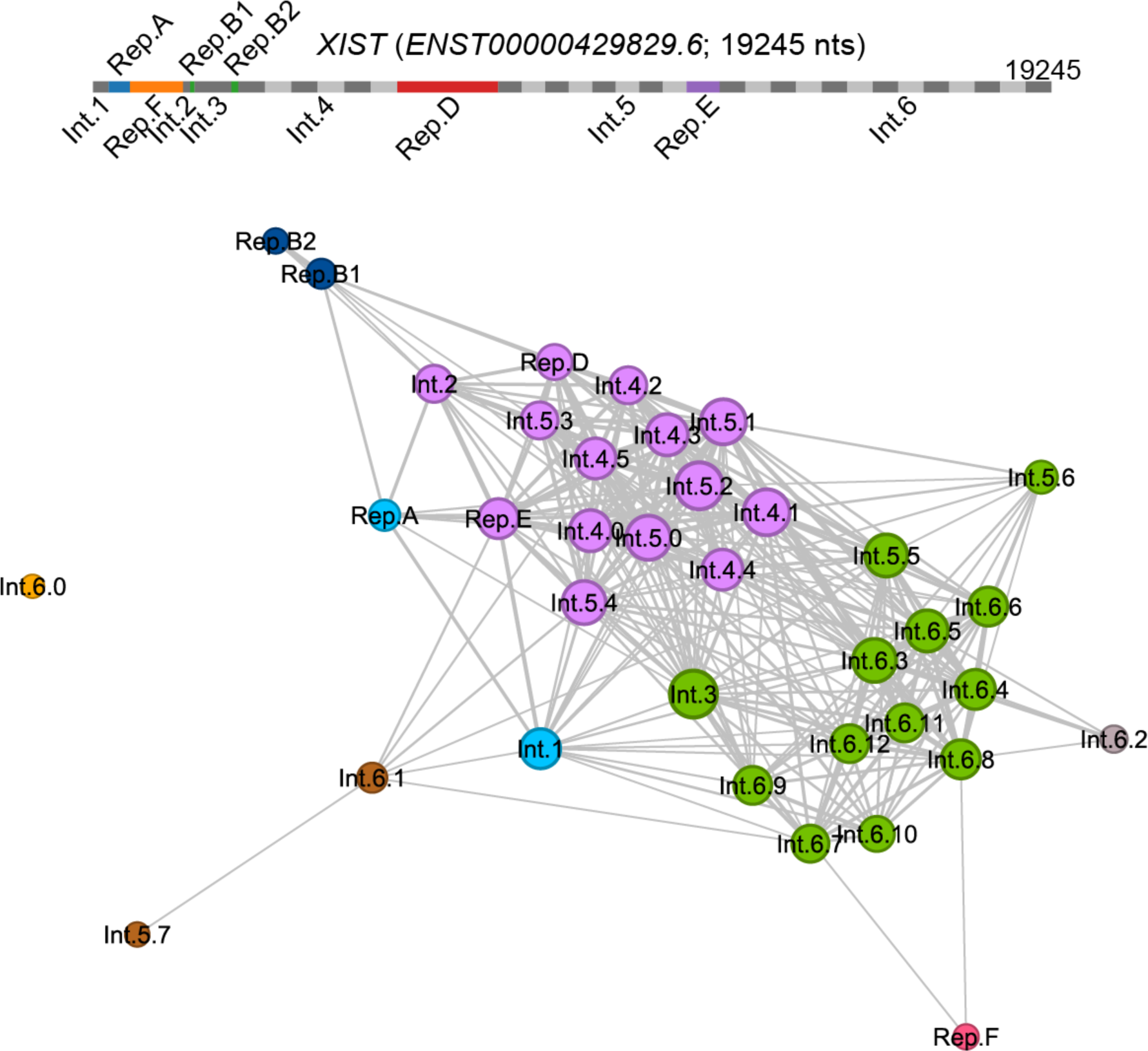
Leiden community graph demonstrating similarity between fragments in the 3′ half of *XIST*. Each circle represents a fragment within *XIST*. Circles are colored by their Leiden community assignments using the RBERVertexPartition function and a Resolution factor of 1.1, and scaled by their number of connections to positive edges. Only those edges that have a Pearson’s r value of greater than 0.15 were counted and displayed here. This image was made using Gephi (Bastian, Heymann et al. 2009).

## Works Cited

Altschul, S. F., W. Gish, W. Miller, E. W. Myers and D. J. Lipman (1990). “Basic local alignment search tool.” J Mol Biol 215(3): 403–410.

Bailey, T. L., J. Johnson, C. E. Grant and W. S. Noble (2015). “The MEME Suite.” Nucleic Acids Res 43(W1): W39–49.

Bastian, M., S. Heymann and M. Jacomy (2009). “Gephi: an open source software for exploring and manipulating networks.” International AAAI Conference on Weblogs and Social Media.

Bray, N. L., H. Pimentel, P. Melsted and L. Pachter (2016). “Near-optimal probabilistic RNA-seq quantification.” Nat Biotechnol 34(5): 525–527.

de Goede, O. M., D. C. Nachun, N. M. Ferraro, M. J. Gloudemans, A. S. Rao, C. Smail, T. Y. Eulalio, F. Aguet, B. Ng, J. Xu, A. N. Barbeira, S. E. Castel, S. Kim-Hellmuth, Y. Park, A. J. Scott, B. J. Strober, G. T. Consortium, C. D. Brown, X. Wen, I. M. Hall, A. Battle, T. Lappalainen, H. K. Im, K. G. Ardlie, S. Mostafavi, T. Quertermous, K. Kirkegaard and S. B. Montgomery (2021). “Population-scale tissue transcriptomics maps long non-coding RNAs to complex disease.” Cell.

Dixon-McDougall, T. and C. J. Brown (2021). “Independent domains for recruitment of PRC1 and PRC2 by human XIST.” PLoS Genet 17(3): e1009123.

Dixon-McDougall, T. and C. J. Brown (2022). “Multiple distinct domains of human XIST are required to coordinate gene silencing and subsequent heterochromatin formation.” Epigenetics Chromatin 15(1): 6.

Dobin, A., C. A. Davis, F. Schlesinger, J. Drenkow, C. Zaleski, S. Jha, P. Batut, M. Chaisson and T. R. Gingeras (2013). “STAR: ultrafast universal RNA-seq aligner.” Bioinformatics 29(1): 15–21.

Dunham, I., A. Kundaje, S. F. Aldred, P. J. Collins, C. A. Davis, F. Doyle, C. B. Epstein, S. Frietze, J. Harrow, R. Kaul, J. Khatun, B. R. Lajoie, S. G. Landt, B. K. Lee, F. Pauli, K. R. Rosenbloom, P. Sabo, A. Safi, A. Sanyal, N. Shoresh, J. M. Simon, L. Song, N. D. Trinklein, R. C. Altshuler, E. Birney, J. B. Brown, C. Cheng, S. Djebali, X. Dong, J. Ernst, T. S. Furey, M. Gerstein, B. Giardine, M. Greven, R. C. Hardison, R. S. Harris, J. Herrero, M. M. Hoffman, S. Iyer, M. Kelllis, P. Kheradpour, T. Lassman, Q. Li, X. Lin, G. K. Marinov, A. Merkel, A. Mortazavi, S. C. Parker, T. E. Reddy, J. Rozowsky, F. Schlesinger, R. E. Thurman, J. Wang, L. D. Ward, T. W. Whitfield, S. P. Wilder, W. Wu, H. S. Xi, K. Y. Yip, J. Zhuang, B. E. Bernstein, E. D. Green, C. Gunter, M. Snyder, M. J. Pazin, R. F. Lowdon, L. A. Dillon, L. B. Adams, C. J. Kelly, J. Zhang, J. R. Wexler, P. J. Good, E. A. Feingold, G. E. Crawford, J. Dekker, L. Elinitski, P. J. Farnham, M. C. Giddings, T. R. Gingeras, R. Guigo, T. J. Hubbard, M. Kellis, W. J. Kent, J. D. Lieb, E. H. Margulies, R. M. Myers, J. A. Starnatoyannopoulos, S. A. Tennebaum, Z. Weng, K. P. White, B. Wold, Y. Yu, J. Wrobel, B. A. Risk, H. P. Gunawardena, H. C. Kuiper, C. W. Maier, L. Xie, X. Chen, T. S. Mikkelsen, S. Gillespie, A. Goren, O. Ram, X. Zhang, L. Wang, R. Issner, M. J. Coyne, T. Durham, M. Ku, T. Truong, M. L. Eaton, A. Dobin, T. Lassmann, A. Tanzer, J. Lagarde, W. Lin, C. Xue, B. A. Williams, C. Zaleski, M. Roder, F. Kokocinski, R. F. Abdelhamid, T. Alioto, I. Antoshechkin, M. T. Baer, P. Batut, I. Bell, K. Bell, S. Chakrabortty, J. Chrast, J. Curado, T. Derrien, J. Drenkow, E. Dumais, J. Dumais, R. Duttagupta, M. Fastuca, K. Fejes-Toth, P. Ferreira, S. Foissac, M. J. Fullwood, H. Gao, D. Gonzalez, A. Gordon, C. Howald, S. Jha, R. Johnson, P. Kapranov, B. King, C. Kingswood, G. Li, O. J. Luo, E. Park, J. B. Preall, K. Presaud, P. Ribeca, D. Robyr, X. Ruan, M. Sammeth, K. S. Sandu, L. Schaeffer, L. H. See, A. Shahab, J. Skancke, A. M. Suzuki, H. Takahashi, H. Tilgner, D. Trout, N. Walters, H. Wang, Y. Hayashizaki, A. Reymond, S. E. Antonarakis, G. J. Hannon, Y. Ruan, P. Carninci, C. A. Sloan, K. Learned, V. S. Malladi, M. C. Wong, G. P. Barber, M. S. Cline, T. R. Dreszer, S. G. Heitner, D. Karolchik, V. M. Kirkup, L. R. Meyer, J. C. Long, M. Maddren, B. J. Raney, L. L. Grasfeder, P. G. Giresi, A. Battenhouse, N. C. Sheffield, K. A. Showers, D. London, A. A. Bhinge, C. Shestak, M. R. Schaner, S. K. Kim, Z. Z. Zhang, P. A. Mieczkowski, J. O. Mieczkowska, Z. Liu, R. M. McDaniell, Y. Ni, N. U. Rashid, M. J. Kim, S. Adar, Z. Zhang, T. Wang, D. Winter, D. Keefe, V. R. Iyer, K. S. Sandhu, M. Zheng, P. Wang, J. Gertz, J. Vielmetter, E. C. Partridge, K. E. Varley, C. Gasper, A. Bansal, S. Pepke, P. Jain, H. Amrhein, K. M. Bowling, M. Anaya, M. K. Cross, M. A. Muratet, K. M. Newberry, K. McCue, A. S. Nesmith, K. I. Fisher-Aylor, B. Pusey, G. DeSalvo, S. L. Parker, S. Balasubramanian, N. S. Davis, S. K. Meadows, T. Eggleston, J. S. Newberry, S. E. Levy, D. M. Absher, W. H. Wong, M. J. Blow, A. Visel, L. A. Pennachio, L. Elnitski, H. M. Petrykowska, A. Abyzov, B. Aken, D. Barrell, G. Barson, A. Berry, A. Bignell, V. Boychenko, G. Bussotti, C. Davidson, G. Despacio-Reyes, M. Diekhans, I. Ezkurdia, A. Frankish, J. Gilbert, J. M. Gonzalez, E. Griffiths, R. Harte, D. A. Hendrix, T. Hunt, I. Jungreis, M. Kay, E. Khurana, J. Leng, M. F. Lin, J. Loveland, Z. Lu, D. Manthravadi, M. Mariotti, J. Mudge, G. Mukherjee, C. Notredame, B. Pei, J. M. Rodriguez, G. Saunders, A. Sboner, S. Searle, C. Sisu, C. Snow, C. Steward, E. Tapanari, M. L. Tress, M. J. van Baren, S. Washieti, L. Wilming, A. Zadissa, Z. Zhengdong, M. Brent, D. Haussler, A. Valencia, A. Raymond, N. Addleman, R. P. Alexander, R. K. Auerbach, K. Bettinger, N. Bhardwaj, A. P. Boyle, A. R. Cao, P. Cayting, A. Charos, Y. Cheng, C. Eastman, G. Euskirchen, J. D. Fleming, F. Grubert, L. Habegger, M. Hariharan, A. Harmanci, S. Iyenger, V. X. Jin, K. J. Karczewski, M. Kasowski, P. Lacroute, H. Lam, N. Larnarre-Vincent, J. Lian, M. Lindahl-Allen, R. Min, B. Miotto, H. Monahan, Z. Moqtaderi, X. J. Mu, H. O’Geen, Z. Ouyang, D. Patacsil, D. Raha, L. Ramirez, B. Reed, M. Shi, T. Slifer, H. Witt, L. Wu, X. Xu, K. K. Yan, X. Yang, K. Struhl, S. M. Weissman, S. A. Tenebaum, L. O. Penalva, S. Karmakar, R. R. Bhanvadia, A. Choudhury, M. Domanus, L. Ma, J. Moran, A. Victorsen, T. Auer, L. Centarin, M. Eichenlaub, F. Gruhl, S. Heerman, B. Hoeckendorf, D. Inoue, T. Kellner, S. Kirchmaier, C. Mueller, R. Reinhardt, L. Schertel, S. Schneider, R. Sinn, B. Wittbrodt, J. Wittbrodt, G. Jain, G. Balasundaram, D. L. Bates, R. Byron, T. K. Canfield, M. J. Diegel, D. Dunn, A. K. Ebersol, T. Frum, K. Garg, E. Gist, R. S. Hansen, L. Boatman, E. Haugen, R. Humbert, A. K. Johnson, E. M. Johnson, T. M. Kutyavin, K. Lee, D. Lotakis, M. T. Maurano, S. J. Neph, F. V. Neri, E. D. Nguyen, H. Qu, A. P. Reynolds, V. Roach, E. Rynes, M. E. Sanchez, R. S. Sandstrom, A. O. Shafer, A. B. Stergachis, S. Thomas, B. Vernot, J. Vierstra, S. Vong, M. A. Weaver, Y. Yan, M. Zhang, J. A. Akey, M. Bender, M. O. Dorschner, M. Groudine, M. J. MacCoss, P. Navas, G. Stamatoyannopoulos, J. A. Stamatoyannopoulos, K. Beal, A. Brazma, P. Flicek, N. Johnson, M. Lukk, N. M. Luscombe, D. Sobral, J. M. Vaquerizas, S. Batzoglou, A. Sidow, N. Hussami, S. Kyriazopoulou-Panagiotopoulou, M. W. Libbrecht, M. A. Schaub, W. Miller, P. J. Bickel, B. Banfai, N. P. Boley, H. Huang, J. J. Li, W. S. Noble, J. A. Bilmes, O. J. Buske, A. O. Sahu, P. V. Kharchenko, P. J. Park, D. Baker, J. Taylor and L. Lochovsky (2012). “An integrated encyclopedia of DNA elements in the human genome.” Nature 489(7414): 57–74.

Frankish, A., S. Carbonell-Sala, M. Diekhans, I. Jungreis, J. E. Loveland, J. M. Mudge, C. Sisu, J. C. Wright, C. Arnan, I. Barnes, A. Banerjee, R. Bennett, A. Berry, A. Bignell, C. Boix, F. Calvet, D. Cerdan-Velez, F. Cunningham, C. Davidson, S. Donaldson, C. Dursun, R. Fatima, S. Giorgetti, C. G. Giron, J. M. Gonzalez, M. Hardy, P. W. Harrison, T. Hourlier, Z. Hollis, T. Hunt, B. James, Y. Jiang, R. Johnson, M. Kay, J. Lagarde, F. J. Martin, L. M. Gomez, S. Nair, P. Ni, F. Pozo, V. Ramalingam, M. Ruffier, B. M. Schmitt, J. M. Schreiber, E. Steed, M. M. Suner, D. Sumathipala, I. Sycheva, B. Uszczynska-Ratajczak, E. Wass, Y. T. Yang, A. Yates, Z. Zafrulla, J. S. Choudhary, M. Gerstein, R. Guigo, T. J. P. Hubbard, M. Kellis, A. Kundaje, B. Paten, M. L. Tress and P. Flicek (2023). “GENCODE: reference annotation for the human and mouse genomes in 2023.” Nucleic Acids Res 51(D1): D942–D949.

Frankish, A., M. Diekhans, I. Jungreis, J. Lagarde, J. E. Loveland, J. M. Mudge, C. Sisu, J. C. Wright, J. Armstrong, I. Barnes, A. Berry, A. Bignell, C. Boix, S. Carbonell Sala, F. Cunningham, T. Di Domenico, S. Donaldson, I. T. Fiddes, C. Garcia Giron, J. M. Gonzalez, T. Grego, M. Hardy, T. Hourlier, K. L. Howe, T. Hunt, O. G. Izuogu, R. Johnson, F. J. Martin, L. Martinez, S. Mohanan, P. Muir, F. C. P. Navarro, A. Parker, B. Pei, F. Pozo, F. C. Riera, M. Ruffier, B. M. Schmitt, E. Stapleton, M. M. Suner, I. Sycheva, B. Uszczynska-Ratajczak, M. Y. Wolf, J. Xu, Y. T. Yang, A. Yates, D. Zerbino, Y. Zhang, J. S. Choudhary, M. Gerstein, R. Guigo, T. J. P. Hubbard, M. Kellis, B. Paten, M. L. Tress and P. Flicek (2021). “Gencode 2021.” Nucleic Acids Res 49(D1): D916–D923.

Hezroni, H., R. Ben-Tov Perry, N. Gil, N. Degani and I. Ulitsky (2020). “Regulation of neuronal commitment in mouse embryonic stem cells by the Reno1/Bahcc1 locus.” EMBO Rep 21(11): e51264.

Kent, W. J., A. S. Zweig, G. Barber, A. S. Hinrichs and D. Karolchik (2010). “BigWig and BigBed: enabling browsing of large distributed datasets.” Bioinformatics 26(17): 2204–2207.

Kirk, J. M., S. O. Kim, K. Inoue, M. J. Smola, D. M. Lee, M. D. Schertzer, J. S. Wooten, A. R. Baker, D. Sprague, D. W. Collins, C. R. Horning, S. Wang, Q. Chen, K. M. Weeks, P. J. Mucha and J. M. Calabrese (2018). “Functional classification of long non-coding RNAs by k-mer content.” Nat Genet 50(10): 1474–1482.

Kirk, J. M., D. Sprague and J. M. Calabrese (2021). “Classification of Long Noncoding RNAs by k-mer Content.” Methods Mol Biol 2254: 41–60.

Levin, J. Z., M. Yassour, X. Adiconis, C. Nusbaum, D. A. Thompson, N. Friedman, A. Gnirke and A. Regev (2010). “Comprehensive comparative analysis of strand-specific RNA sequencing methods.” Nat Methods 7(9): 709–715.

Li, H., B. Handsaker, A. Wysoker, T. Fennell, J. Ruan, N. Homer, G. Marth, G. Abecasis, R. Durbin and S. Genome Project Data Processing (2009). “The Sequence Alignment/Map format and SAMtools.” Bioinformatics 25(16): 2078–2079.

Luo, Y., B. C. Hitz, I. Gabdank, J. A. Hilton, M. S. Kagda, B. Lam, Z. Myers, P. Sud, J. Jou, K. Lin, U. K. Baymuradov, K. Graham, C. Litton, S. R. Miyasato, J. S. Strattan, O. Jolanki, J. W. Lee, F. Y. Tanaka, P. Adenekan, E. O’Neill and J. M. Cherry (2020). “New developments on the Encyclopedia of DNA Elements (ENCODE) data portal.” Nucleic Acids Res 48(D1): D882–D889.

Mattick, J. S., P. P. Amaral, P. Carninci, S. Carpenter, H. Y. Chang, L. L. Chen, R. Chen, C. Dean, M. E. Dinger, K. A. Fitzgerald, T. R. Gingeras, M. Guttman, T. Hirose, M. Huarte, R. Johnson, C. Kanduri, P. Kapranov, J. B. Lawrence, J. T. Lee, J. T. Mendell, T. R. Mercer, K. J. Moore, S. Nakagawa, J. L. Rinn, D. L. Spector, I. Ulitsky, Y. Wan, J. E. Wilusz and M. Wu (2023). “Long non-coding RNAs: definitions, functions, challenges and recommendations.” Nat Rev Mol Cell Biol 24(6): 430–447.

Nassar, L. R., G. P. Barber, A. Benet-Pages, J. Casper, H. Clawson, M. Diekhans, C. Fischer, J. N. Gonzalez, A. S. Hinrichs, B. T. Lee, C. M. Lee, P. Muthuraman, B. Nguy, T. Pereira, P. Nejad, G. Perez, B. J. Raney, D. Schmelter, M. L. Speir, B. D. Wick, A. S. Zweig, D. Haussler, R. M. Kuhn, M. Haeussler and W. J. Kent (2023). “The UCSC Genome Browser database: 2023 update.” Nucleic Acids Res 51(D1): D1188–D1195.

Obuse, C. and T. Hirose (2023). “Functional domains of nuclear long noncoding RNAs: Insights into gene regulation and intracellular architecture.” Curr Opin Cell Biol 85: 102250.

Pandya-Jones, A., Y. Markaki, J. Serizay, T. Chitiashvili, W. R. Mancia Leon, A. Damianov, C. Chronis, B. Papp, C. K. Chen, R. McKee, X. J. Wang, A. Chau, S. Sabri, H. Leonhardt, S. Zheng, M. Guttman, D. L. Black and K. Plath (2020). “A protein assembly mediates Xist localization and gene silencing.” Nature 587(7832): 145–151.

Quinlan, A. R. and I. M. Hall (2010). “BEDTools: a flexible suite of utilities for comparing genomic features.” Bioinformatics 26(6): 841–842.

Quinodoz, S. A., J. W. Jachowicz, P. Bhat, N. Ollikainen, A. K. Banerjee, I. N. Goronzy, M. R. Blanco, P. Chovanec, A. Chow, Y. Markaki, J. Thai, K. Plath and M. Guttman (2021). “RNA promotes the formation of spatial compartments in the nucleus.” Cell 184(23): 5775–5790 e5730.

Raney, B. J., T. R. Dreszer, G. P. Barber, H. Clawson, P. A. Fujita, T. Wang, N. Nguyen, B. Paten, A. S. Zweig, D. Karolchik and W. J. Kent (2014). “Track data hubs enable visualization of user-defined genome-wide annotations on the UCSC Genome Browser.” Bioinformatics 30(7): 1003–1005.

Schertzer, M. D., K. C. A. Braceros, J. Starmer, R. E. Cherney, D. M. Lee, G. Salazar, M. Justice, S. R. Bischoff, D. O. Cowley, P. Ariel, M. J. Zylka, J. M. Dowen, T. Magnuson and J. M. Calabrese (2019). “lncRNA-Induced Spread of Polycomb Controlled by Genome Architecture, RNA Abundance, and CpG Island DNA.” Mol Cell 75(3): 523–537 e510.

Sprague, D., S. A. Waters, J. M. Kirk, J. R. Wang, P. B. Samollow, P. D. Waters and J. M. Calabrese (2019). “Nonlinear sequence similarity between the Xist and Rsx long noncoding RNAs suggests shared functions of tandem repeat domains.” RNA 25(8): 1004–1019.

Sunwoo, H., D. Colognori, J. E. Froberg, Y. Jeon and J. T. Lee (2017). “Repeat E anchors Xist RNA to the inactive X chromosomal compartment through CDKN1A-interacting protein (CIZ1).” Proc Natl Acad Sci U S A 114(40): 10654–10659.

Trotman, J. B., K. C. A. Braceros, R. E. Cherney, M. M. Murvin and J. M. Calabrese (2021). “The control of polycomb repressive complexes by long noncoding RNAs.” Wiley Interdiscip Rev RNA: e1657.

van Dijk, M., H. K. Thulluru, J. Mulders, O. J. Michel, A. Poutsma, S. Windhorst, G. Kleiverda, D. Sie, A. M. Lachmeijer and C. B. Oudejans (2012). “HELLP babies link a novel lincRNA to the trophoblast cell cycle.” J Clin Invest 122(11): 4003–4011.

Van Nostrand, E. L., P. Freese, G. A. Pratt, X. Wang, X. Wei, R. Xiao, S. M. Blue, J. Y. Chen, N. A. L. Cody, D. Dominguez, S. Olson, B. Sundararaman, L. Zhan, C. Bazile, L. P. B. Bouvrette, J. Bergalet, M. O. Duff, K. E. Garcia, C. Gelboin-Burkhart, M. Hochman, N. J. Lambert, H. Li, M. P. McGurk, T. B. Nguyen, T. Palden, I. Rabano, S. Sathe, R. Stanton, A. Su, R. Wang, B. A. Yee, B. Zhou, A. L. Louie, S. Aigner, X. D. Fu, E. Lecuyer, C. B. Burge, B. R. Graveley and G. W. Yeo (2020). “A large-scale binding and functional map of human RNA-binding proteins.” Nature 583(7818): 711–719.

Van Nostrand, E. L., G. A. Pratt, A. A. Shishkin, C. Gelboin-Burkhart, M. Y. Fang, B. Sundararaman, S. M. Blue, T. B. Nguyen, C. Surka, K. Elkins, R. Stanton, F. Rigo, M. Guttman and G. W. Yeo (2016). “Robust transcriptome-wide discovery of RNA-binding protein binding sites with enhanced CLIP (eCLIP).” Nat Methods 13(6): 508–514.

Yamada, N., Y. Hasegawa, M. Yue, T. Hamada, S. Nakagawa and Y. Ogawa (2015). “Xist Exon 7 Contributes to the Stable Localization of Xist RNA on the Inactive X-Chromosome.” PLoS Genet 11(8): e1005430.

Yue, M., A. Ogawa, N. Yamada, J. L. Charles Richard, A. Barski and Y. Ogawa (2017). “Xist RNA repeat E is essential for ASH2L recruitment to the inactive X and regulates histone modifications and escape gene expression.” PLoS Genet 13(7): e1006890.

Zhu, X., M. Du, H. Gu, R. Wu, M. Gao, H. Xu, J. Tang, M. Li, X. Liu and X. Zhong (2023). “Integrated analysis of lncRNA and mRNA expression profiles in patients with unexplained recurrent spontaneous abortion.” Am J Reprod Immunol 89(6): e13691.

